# Moremi Bio Agent: Leveraging Agentic Large Language Model for the Discovery of Broad-Spectrum Antibiotics for Enterobacteriaceae

**DOI:** 10.1101/2025.08.21.671656

**Authors:** Gertrude Hattoh, Jeremiah Ayensu, Nyarko Prince Ofori, Solomon Eshun, Joshua Ntow Opare-Boateng, Osman Tanko, Darlington Akogo

## Abstract

Antimicrobial resistance (AMR) is a pressing global health crisis, exacerbated by a stagnating antibiotic discovery pipeline and the emergence of multidrug-resistant pathogens such as *Klebsiella pneumoniae*. We hypothesize that dual-target strategies may offer a more robust means to overcome AMR by reducing the likelihood of resistance development compared to single-target approaches. In response, we leveraged Moremi Bio Agent, an agentic large language model (LLM) for the autonomous design, in silico validation, and prioritization of broad-spectrum antibiotics targeting Enterobacteriaceae. Using our proposed dual-target strategy, we generated and evaluated 1,002 candidate molecules predicted to simultaneously inhibit the FabI enzyme and the AcrAB–TolC efflux pump—two key resistance mechanisms in Gram-negative bacteria. Our fully autonomous pipeline integrated compound generation, molecular docking, pharmacodynamics/pharmacokinetic predictions & toxicity profiling, and molecule-ranking based on ADMET and drug-likeness properties. Out of 1,002 molecules generated, 774 passed preliminary ADMET benchmarks, with majority of the 60 top-performing candidates (score *≥* 0.8) showing favorable drug-likeness, minimal toxicity. 391 of the compounds exhibited moderate binding interaction to both targets. This study demonstrates the feasibility of AI-driven antibiotic discovery and lays the foundation for future experimental validation to address AMR.

## 1 Introduction

Antimicrobial Resistance (AMR) is a significant global health threat, recognized by the World Health Organization (WHO) as one of the top global public health threats facing humanity World Health Organization (2023). This has been a critical challenge to healthcare and antimicrobial research, largely due to overuse and misuse of antimicrobial drugs Salam et al. (2023). The Global Research on Antimicrobial Resistance, GRAM reports that each year, AMR causes significant deaths worldwide; approximately 1.27 million deaths were recorded in 2019 Kim et al. (2022) and a projected economic burden of $100 trillion USD by 2050 if no effective new antibiotics are developed O’neill (2014).

The global fight against AMR faces significant challenges due to a stagnant antibiotic pipeline Årdal et al. (2019). Despite urgent needs, there is a critical shortfall in new antibiotics reaching clinical development. This is largely attributed to high research and development (R&D) costs Schlander et al. (2021), which deter investment in antibiotic innovation. The current pipeline is described as inadequate to address the mounting threat of antibiotic resistance, leaving patients vulnerable to increasingly resistant bacterial infections. Consequently, the limited discovery of novel antibiotics has significantly hindered efforts to combat AMR Laxminarayan et al. (2013), allowing drug-resistant pathogens to proliferate unchecked. This necessitates the need for new therapeutic approaches Vaja et al. (2024).

Amidst these challenges, Artificial Intelligence (AI), particularly Large Language Models (LLMs), has emerged as a transformative force in biotechnology, revolutionizing drug discovery, protein engineering, and antibody development Xiang et al. (2024). In the context of antimicrobial resistance (AMR), AI-driven approaches have shown substantial promise across key domains, including discovery and design, diagnostic support, and data integration—each contributing to accelerated progress against AMR Yoo et al. (2024). Notable advancements include the use of generative models to design antibiotics targeting *Acinetobacter baumannii* Swanson et al. (2024a), the development of novel antimicrobial peptides Wang et al. (2024), molecular optimization, and the acceleration of early-stage drug discoveryTorres et al. (2024), Delmas et al. (2025).

Amidst global efforts to tackle antimicrobial resistance (AMR), Minohealth AI Labs is leading innovation in Africa and beyond. Our AI model, Moremi Bio Agent—already proven in autonomously designing and validating novel antibodies against malaria and small molecules for diverse diseases Akogo et al. (2025), MinohealthAILabs (2025) —has now been deployed to generate an initial library of 1,002 novel broad-spectrum antibiotic candidates targeting Enterobacteriaceae, using Klebsiella species as the starting point. This foundational effort aims to address critical gaps in antibiotic discovery research. Our approach not only accelerates the rapid generation of novel antibiotic scaffolds but also explores the potential of a dual-target strategy—a hypothesis we propose may offer a more robust means to overcome AMR by reducing the likelihood of resistance development. Through comprehensive in silico high-throughput design, validation, and iterative optimization, we will refine and prioritize the most promising candidates, advancing them through successive stages, from computational refinement to possible design enhancements and experimental testing. This initiative sets the stage for a scalable and high-throughput framework designed to meet the AMR challenge where traditional methods have struggled.

## 2 Literature Review

### 2.1 Background

Antimicrobial resistance (AMR) is an escalating global health crisis, contributing to substantial morbidity, mortality, and economic burden worldwide 201 (2019). Infections caused by multidrug-resistant pathogens—especially members of the Enterobacteriaceae family such as *Klebsiella* spp., pose a severe threat to public health Podschun and Ullmann (1998). *Klebsiella pneumoniae*, in particular, is notorious for its ability to acquire resistance mechanisms, rendering many conventional antibiotics ineffective and complicating treatment regimens for critically ill patients Li et al. (2023, 2024).

Traditional antibiotic discovery methods, including high-throughput screening and conventional computational approaches, have been hampered by extensive timeframes, high costs, and limited success rates. These methods often fail to explore the vast chemical space comprehensively, thereby missing novel chemical scaffolds with unique multi-target mechanisms that are critical for circumventing bacterial resistance Terekhov et al. (2018), Silver (2011). Recent landmark discoveries—such as the identification of Halicin through artificial intelligence—have demonstrated the potential of machine learning (ML) techniques to uncover new antibiotics. However, these advancements also underscore the ongoing need for innovative approaches to rapidly identify and validate antibiotic candidates with novel mechanisms of action, which can address the growing challenge of antimicrobial resistance. Possibly by integrating multi-target design strategies into their generative frameworks, LLMs can explore vast chemical spaces and propose novel antibiotic structures in a matter of minutes—a stark contrast to the prolonged timelines of traditional experimental methods.

In this study, we present an innovative approach that harnesses LLMs to accelerate the discovery of novel broad-spectrum antibiotics specifically targeted against members of the Enterobacteriacea starting with *Klebsiella* spp. Our method proposes a dual-target strategy to design candidate molecules that are predicted to interact with two essential bacterial pathways Gray and Wenzel (2020). These candidate molecules are then subjected to rigorous in silico assessments, including evaluations of their Absorption, Distribution, Metabolism, Excretion and Toxicity (ADMET) properties, stability, and binding affinities with proposed bacterial targets. This integrated computational pipeline not only significantly reduces the time and cost associated with antibiotic development but also enhances the likelihood of identifying candidates with robust efficacy against members of Enterobacteriaceae.

### 2.2 The Global Urgency for Gram-Negative Antibiotics

According to the WHO Priority Pathogens List, Gram-negative bacteria represent one of the foremost challenges in combating antimicrobial resistance Moja et al. (2017). In combating AMR, it is essential to prioritize Gram-negative bacteria since they exhibit greater resistance than their Gram-positive counterparts—owing to their inherent defense mechanisms, intricate cell architecture, and the increasing burden of hospital-acquired infections Melander et al. (2022).

The WHO’s priority list highlights critical Gram negative pathogens including the family Enterobacteriaceae as those among the ‘critical’ class Moja et al. (2017). Many reports indicate that resistance to key antibiotics such as third-generation cephalosporins and carbapenems is rising within this family. For instance, WHO data and regional surveillance efforts have documented that in some countries, up to 42% of E. coli isolates are resistant to third-generation cephalosporins, while similar high resistance levels are seen among *K. pneumoniae*. These trends contribute heavily to the morbidity and mortality from bloodstream and urinary tract infections, among other conditions G et al. (2022).

Gram-negative pathogens owe much of their antimicrobial resistance to their complex cell envelope architecture: a dense lipopolysaccharide (LPS) outer membrane that restricts drug entry, porin modifications that further limit influx, robust efflux pump systems that expel antibiotics, and a suite of *β*-lactamases and other enzymes that degrade drugs before they can act. These intertwined barriers underscore the urgent need for innovative therapeutic strategies that can breach or bypass them. While approaches such as phage therapy and immunomodulation are gaining traction Bhandari and Suresh (2022), World Health Organization (2023), Alipour-Khezri et al. (2024), antibiotics remain indispensable, even as their efficacy wanes. To invigorate the antibiotic arsenal, we propose coupling LLM-powered exploration of vast chemical space with a polypharmacology paradigm to design single molecules that engage two bacterial targets. We hypothesize that such multi-target (dual-action) agents will significantly lower the chance of resistance emergence by striking two critical pathways at once Feng et al. (2023). Embedding this concept within a structured pipeline—spanning in silico discovery, rigorous computational validation, and iterative optimization—offers a path toward next-generation antibiotics capable of overcoming Gram-negative defenses where traditional monotherapies have repeatedly fallen short Khabthani et al. (2021).

### 2.3 Significance of Targeting Enterobacteriaceae and Klebsiella species

Enterobacteriaceae represent the largest and most diverse family of Gram-negative bacteria Oliveira and Reygaert (2023). Among these, *Escherichia coli* and *Klebsiella pneumoniae* are consistently identified as the leading pathogens in Hospital-acquired infections (HIAs) Irek et al. (2018), Saleem et al. (2019). The 2024 WHO Bacterial Priority Pathogens List (BPPL) classifies carbapenemresistant Enterobacterales—including carbapenem-resistant *K. pneumoniae*—in its highest priority tier, reflecting their urgent need for novel therapeutics Sati et al. (2025).

*Klebsiella* spp. are ubiquitous and cause a wide range of infections Bengoechea and Pessoa (2018) which includes bloodstream infections, urinary tract infections, pneumonia and other life threatening conditions Xu et al. (2018), Gonzalez-Ferrer et al. (2021), Chang et al. (2021). The pathogenic success of *Klebsiella* spp. and related Enterobacteriaceae stems from an arsenal of resistance mechanisms: production of *β* –lactamases and carbapenemases that inactivate antibiotics, modification of drug targets, loss or mutation of outer-membrane porins, upregulation of efflux pumps, and formation of biofilms that impede antimicrobial penetration Paczosa and Mecsas (2016), Wang et al. (2020).

*Klebsiella* spp, more specially *K. pneumoniae* is an opportunistic pathogen, which causes around 6-10% of nosocomial infections worldwide Navon-Venezia et al. (2017), Asri et al. (2021), Shah et al. (2025). Its role as a major public health concern is heightened by its ability to rapidly develop resistance to multiple antibiotic classes including carbapenem 201 (2019), placing it at the top of the World Health Organization’s 2024 Bacterial Priority Pathogens List. In the United States, the CDC’s 2019 report on Antibiotic Resistance Threats specifically lists Klebsiella as one of the most pressing concerns for antibiotic resistance 201 (2019). *K. pneumoniae* is recognized as an important multidrug-resistant (MDR) pathogen, with increasing prevalence of resistance to first-line and last-resort antibiotics complicating clinical therapy Li et al. (2022, 2023).

*Klebsiella* spp provides a physiologically relevant model for screening compounds against traits common to Enterobacteriaceae; outer-membrane permeability barriers, resistance, virulence and *β*-lactamase activity Li et al. (2025). Also, using Klebsiella spp. as a starting point aligns with global research initiatives (E.g Grand Challenges in Gram-negative Antibiotic Discovery) which designates Klebsiella spp. for early-phase drug discovery to foster collaboration and accelerate the development of broad-spectrum agents against Enterobacteriaceae.^1^ Therefore, a thorough understanding of Klebsiella resistance and pathogenicity is critical for guiding the discovery of novel antibiotics with activity across the Enterobacteriaceae family and for addressing the global antimicrobial resistance crisis.

### 2.4 Fabl and AcrAB-TolC efflux pump

Our goal is to leverage Moremi Bio to autonomously generate and in silico validate one thousand and two (1,002) novel compounds; adopting a polypharmacology strategy, these candidates were designed to inhibit the FabI enzyme while simultaneously blocking the AcrAB-TolC pump in a high-throughput manner. FaBI are enoyl-ACP (Acyl Carrier Protein) reductases critical in the pathway II fatty acid biosynthesis (FAS II) in most bacteria Lu and Tonge (2008). Fatty acids play essential roles in the bacteria from use for membrane synthesis essential for their survival through to contribution of virulence and pathogenesis Waters and Eijkelkamp (2024). FabI occupies a pivotal, rate-determining position in the bacterial FAS II pathway: as the terminal enoyl-ACP reductase, it leverages a highly favorable reduction to pull the elongation cycle to completion and regulate flux into membrane phospholipid assembly Parsons and Rock (2013). FaBI has been considered a significant therapeutic target for antibacterial drug discovery, highly conserved components in many pathogens and are non-homologous to human targets; targeting the Fabl enzyme not only inhibits bacterial infections but also leads to a broad spectrum antimicrobial activity Rana et al. (2020). While bacteria can incorporate extracellular fatty acids, exogenous lipids fail to fully compensate for the essential FAS-II derived intermediates and can not bypass the need for de novo fatty acid synthesis; ultimately blocking fatty synthesis remains critical and an effective antimicrobial strategy Yao et al. (2015), Rana et al. (2020), Biswas et al. (2023). [Fabl structure from the PDB database with PDB 8JTP].

AcrAB-TolC efflux pump is a tripartite complex composed of the AcrA, AcrB and TolC proteins of the Resistance-nodulation-division (RND)-family efflux pumps; a major mechanism by which Klebsiella resists multiple classes of antibiotics Wand et al. (2022). TolC penetrates the outer membrane bacteria lipid bilayer, AcrA is positioned within the layer and forms the bridge between the AcrB on the inner to the TolC on the outer cell membrane Wang et al. (2017), Chen et al. (2020) [Structure from the PDB Database with id = 5O66]. The AcrAB-TolC efflux pump has various roles which includes multidrug resistance, biofilm formation and virulence, and stress adaptation Padilla et al. (2009), Jang (2023).

Targeting both Fabl and the AcrAB-TolC efflux pump simultaneously presents a promising polypharmacological strategy to combat AMR. Fabl inhibition disrupts bacterial membrane biosynthesis by blocking the enoyl-ACP reductase step in fatty acid synthesis Mehboob et al. (2012) and AcrAB-TolC inhibition prevents antibiotics efflux hence increasing the intracellular drug concentrations Alenazy (2022); which intend create a lethal accumulation effect.

## 3 Methodology

This project employed the use of LLM-based approach to mine the literature, identify relevant antibacterial targets, and designed novel antibiotics against Klebsiella spp. Using our model, Moremi Bio, we adopted a polypharmacology strategy to autonomously generate and in silico evaluate one thousand and two (1,002) novel antibiotic candidates. These candidates were designed to inhibit the FabI enzyme while simultaneously blocking the AcrAB-TolC efflux pump in a high-throughput manner.

The compound generation and validation pipeline, which is the pre-developed agentic model capable of autonomously executing these tasks without human in-the-loop was leveraged. Moremi Bio Agent integrated with advanced external industry-standard tools to validate generated antibiotics. The validation process involved molecular docking studies to assess binding affinity with the targets and in silico predictions of the Pharmacodynamics/Pharmacokinetics (PD/PK). The assessment also evaluated toxicity (LD_50_), off-target interactions (hERG), and stability to ensure a comprehensive evaluation of the antibiotic candidates. A ranking system in the pipeline assigned ranks to all molecules, facilitating the easy identification of lead compounds based on their ADMET properties and other relevant properties [See Appendix A]. An overview of the method is presented in an image below, figure 1.

**Figure 1:**
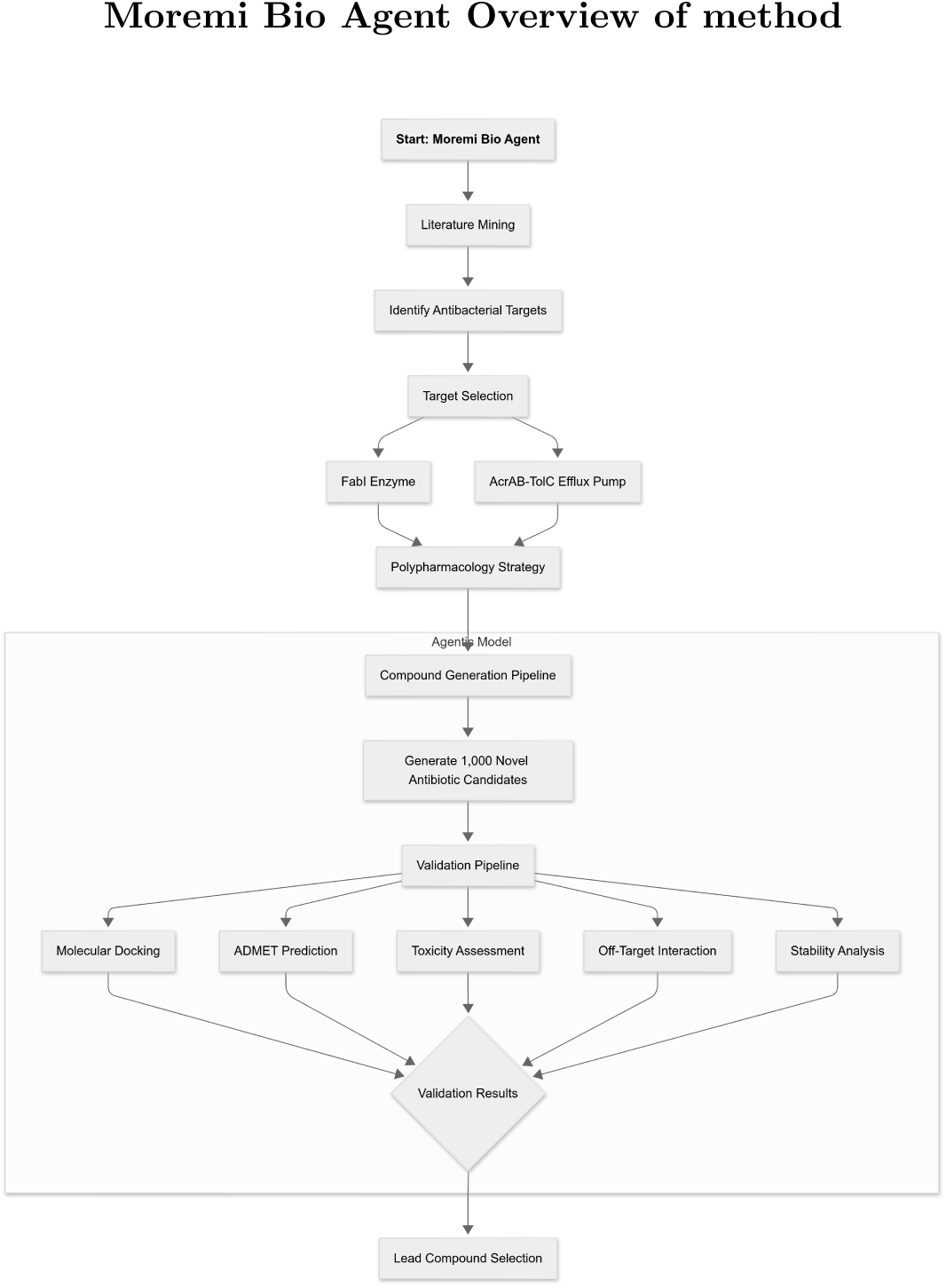
Overview of Moremi Bio Agent method.

A framework of the computational design, agentic pipeline with relevant components and verification of novel compounds.

## 4 Results

This research applied an AI-driven approach to swiftly design unique antibiotic molecules with dual-target inhibition to Fabl and AcrAB-TolC efflux pump, with validated PD/PK properties, and accessed on several critical criteria which includes Binding affinity assessment; providing a promising foundation for future experimental validation against AMR pathogens. Moremi Bio generated a total of 1002 molecules of which 228 failed on several benchmarks. Out of the 774 candidate molecules that passed preliminary benchmarks, 545 (approximately 70.4%) had molecular weights below 700 Da.

### 4.1 PD/PK Properties

Performing in silico assessment of relevant drug-likeness of generated molecules significantly raises the probability in identifying leads and areas for further optimization before further in vitro-in vivo studies Dhoble et al. (2025). PD/PK of molecules were assessed using the ADMETAI Swanson et al. (2024b); these included Absorption, Distribution, Metabolism, Excretion, and Toxicity (ADMET) predictions. Several metrics are inclusive to obtain each category of the ADMET prediction of the novel compounds as predicted through the ADMETAI [See Appendix A]. The figures below summarizes the descriptive and distributive analysis of the generated compounds. This presents a broad overview to the total landscape distribution of all generated molecules across the ADMET properties.

Each subplot shows a histogram overlaid with a kernel density estimate (KDE), illustrating both frequency counts and the underlying distribution shape. The model predicted high absorption (scores *≥* 0.5) for 72% of candidates. The distribution score (subplot2) shows a prominent peak around 0.45 indicating that nearly half of the compounds have moderate tissue distribution and a gentle right shows that fewer molecules achieved scores *≥* 0.8. The Metabolic scores show a pronounced right skew revealing that majority of the compounds were metabolically stable. The Excretion score unlike other metrics shows a bimodal distribution. The toxicity distribution score, based on predicted hERG inhibition probabilities, indicates that a substantial proportion of the compounds exhibit low predicted hERG inhibition (i.e., low probability of cardiotoxicity), reflected in high toxicity safety scores. This suggests that many of the generated molecules are likely to have a favorable cardiac safety profile, with relatively few exhibiting moderate-to-high risk for cardiotoxic effects.

Molecules, each color-coded by its overall score (composite ADMET scores (0-1)) integrating each property listed in A. Those with similar ADMET and drug-likeness profiles cluster together spatially.

Regions with lighter colors (higher scores) suggest clusters of molecules with optimal profiles, potentially highlighting promising drug candidates. Conversely, darker regions represent molecules with less desirable properties. This prediction gives an indication that the overall scores of the molecules somewhat correlates with the structural property hence translating into the molecules’ performance.

A high-resolution comparison of how our novel antibiotic candidates distribute across key ADMET properties offer deep insight into their developability profiles. By juxtaposing summary-statistical box plots with full-distribution violin plots, we can both quantify central tendencies and visualize the underlying density shapes.

### 4.2 Absorption

One of the key determinants of oral efficacy for broad-spectrum antibiotics is their ability to be absorbed efficiently through the gut epithelium; which is mostly evaluated in vitro by measuring permeability across cell-based or artificial membrane models (Wang et al., 2016), predicting Human Intestinal Absorption (HIA) along with assessing interactions with P-glycoprotein (Pgp) Kus et al. (2023) is essential early in silico assessment. Caco-2 (Cancer coli) cell line 2 were predicted to range from *−*7.578 *×* log(10*^−^*^6^) cm/s and *−*3.800 *×* log(10*^−^*^6^) cm/s, which suggest very high absorption in the gut epithelium Kus et al. (2023) for *−*3.800 *×* log(10*^−^*^6^) cm/s (i.e., 158 *×* 10*^−^*^6^ cm/s). HIA reached 0.999 (99.9%) exceeding the 80% threshold for high bioavailability (Yan et al., 2008). Parallel Artificial Membrane Permeability Assay (PAMPA) values which indicate the ability of the molecule to diffuse an artificial membrane were as high as 99.7%. These are indicative that the optimal absorption of the generated molecules can possibly reach their therapeutic concentrations in the plasma and significantly raises the probability that they will achieve the systemic exposures required for broad-spectrum activity.

### 4.3 Distribution

Approximately, sixty-three percent (*≈*63.00%) of the molecules had optimal Plasma Protein Bound (PPB) values (i.e., 50-90% PPB)Watanabe et al. (2018). This means an adequate quantity of unbound compounds exist in the plasma to exert antimicrobial effect.

### 4.4 Metabolism

A minimal interaction with the cytochrome p450 (CYP) enzymes is optimal. Predicted probability of cytochrome inhibition measured for some major CYP (Cyp2C9, Cyp2D6 and Cyp3A4) Arakawa et al. (2023) enzymes ranged from 9.427e-05 to 0.980, 2.295e-04 to 0.922 and 7.871e-06 to 0.993 respectively. CYP2C9 inhibition probabilities up to 0.98 indicate high risk for drug–drug interactions in candidates with hepatic metabolism pathways Boiko et al. (2020). The top-10 ranked candidates showed minimal interaction with the cytochrome p450 enzymes, the mean of the min and max inhibition values across the three CYP inhibition values ranged from o.0016 to 0.1406, suggesting an acceptable range which translates to reduced hepatic elimination, longer in vivo half-lives, and potentially less frequent dosing in humans.

### 4.5 Excretion

Approximately, 37.60% of molecules exhibited Clearance(Cl) less than 5 *µ*L/min/10^6^ cells (*<* 15 mL min*^−^*^1^ kg*^−^*^1^); the large low-Cl fraction suggests that several compounds may achieve sustained plasma exposure ideal for good dosing regimens. The highest clearance rate recorded was 28.462 *µ*L/min/10*^−^*^6^ cells S-loczyńska et al. (2019).

Half-life (*t*_1_*_/_*_2_) inversely correlates with clearance (Cl), as defined by *t*_1_*_/_*_2_ = 0.693×Vd/Cl, demonstrating that half-life is directly proportional to volume of distribution and inversely proportional to clearance Grogan and Preuss (2023). The observed negative correlation in figure 5 aligns with established pharmacokinetic theory. Compounds with elevated clearance demonstrate shorter half-lives, potentially necessitating more frequent dosing regimens. Conversely, lower clearance values typically yield extended half-lives, which may facilitate less frequent administration schedules and more sustained therapeutic plasma concentrations.

**Figure 2:**
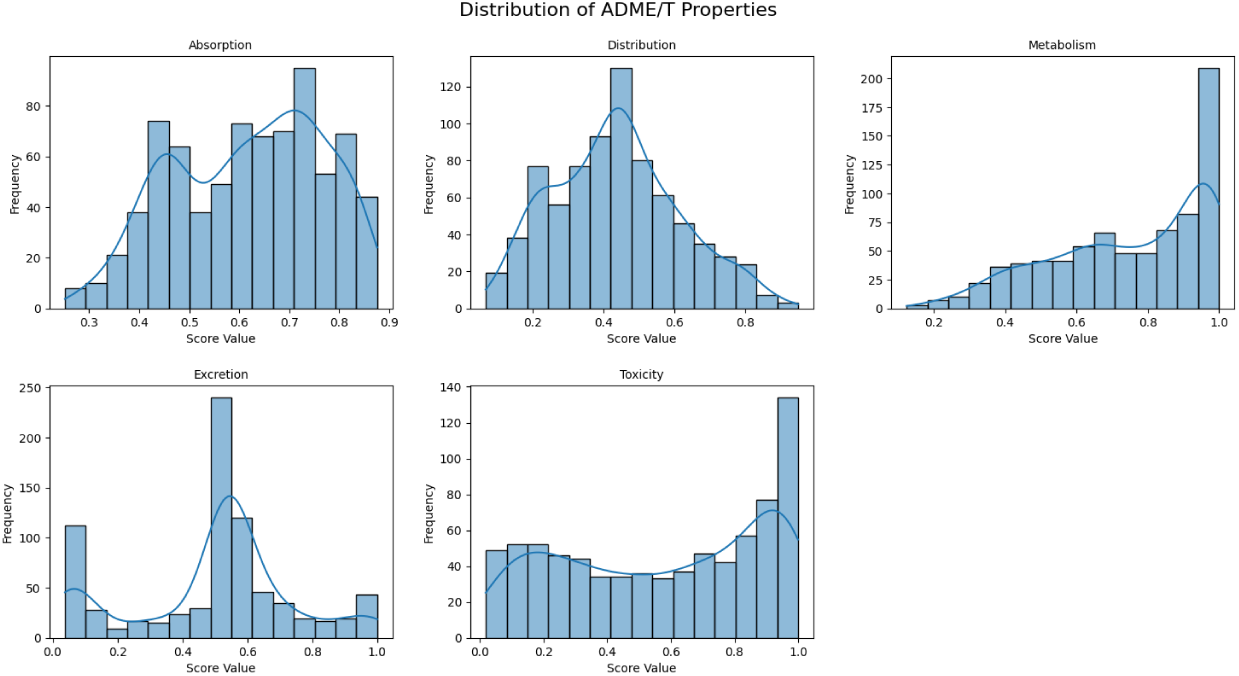
Distribution of compounds profiled access the five ADMET metrics.

**Figure 3:**
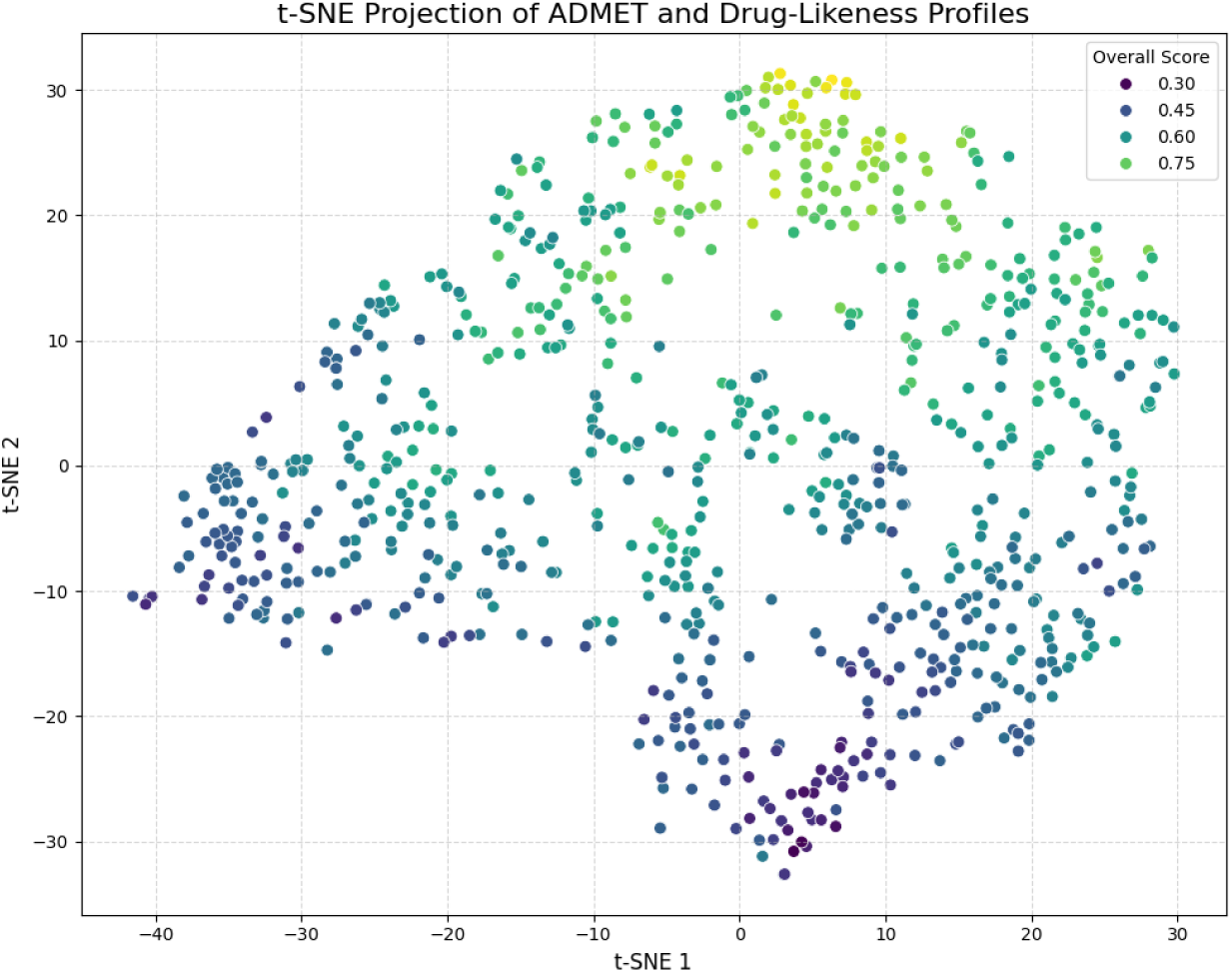
t-SNE projection of ADMET properties and drug-likeness profiles.

**Figure 4:**
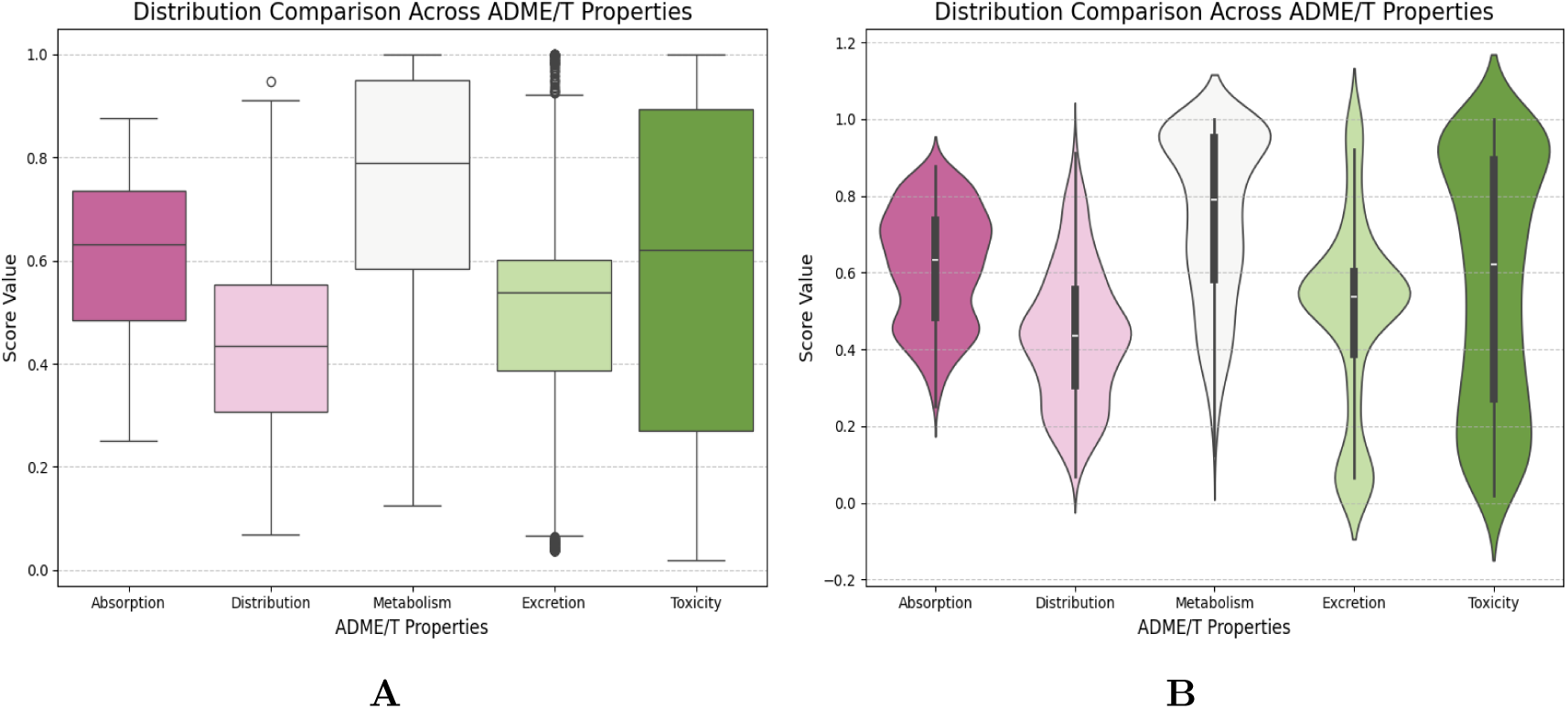
Distribution comparison across ADMET properties. **A**: Box and whisker plot of distribution across each ADMET property. **B**: Violin plot showing comparison across ADMET properties. From the distribution in **A** and **B**, most candidates excel at Metabolism and equally exhibit good Absorption, Distribution, Excretion, and Toxicity.

**Figure 5:**
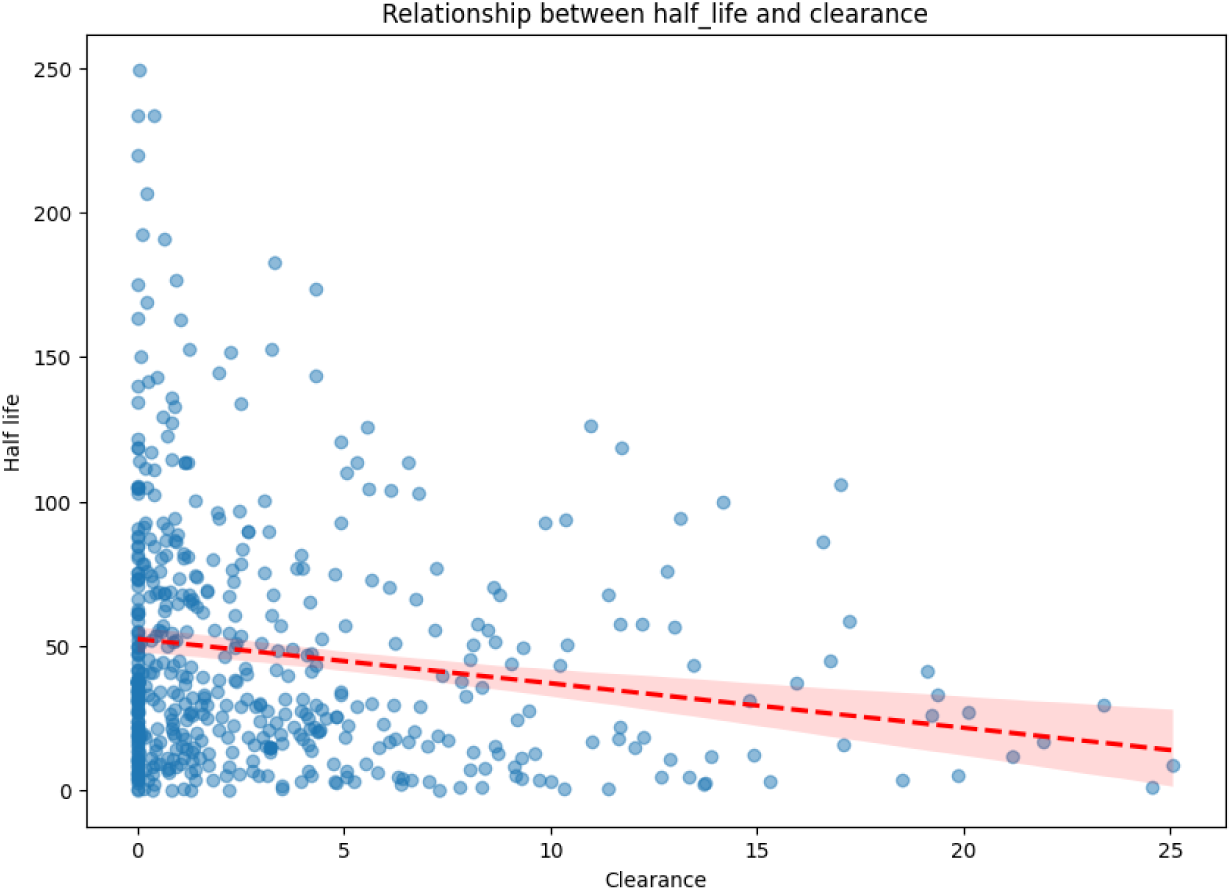
Relationship between half-life and clearance.

The substantial data scatter observed suggest significant variability in the clearance-half-life relationship.

### 4.6 Toxicity and off target interaction

Lethal Dose 50 (LD_50_) and the human Ether-ía-go-go-Related Gene (hERG) inhibition were measured to define the potential toxicity and off-target interaction of the generated molecules; as early assessment of toxicity through in silico means is desirable to ascertain the candidacy for the various generated molecules. hERG is a critical indicator of cardiotoxicity as it encodes the Kv11.1 protein responsible for electrical activity in the heart Vandenberg et al. (2012). The min and max probability of herG inhibition values ranged from 5.393 *×* 10*^−^*^4^ to 9.813 *×* 10*^−^*^1^. This gives an indication that some molecules have high potential to cause cardiac side effects Garrido et al. (2020).Yet, the top-ten ranked molecules 1.799 *×* 10*^−^*^3^ and 3.429 *×* 10*^−^*^1^ (values are closer to zero) indicating a very low probability for hERG inhibition hence suggesting little to no potential cardiac side effects. LD_50_ values range from 1.213 to 4.593 in log(1/(mol/kg)). Table 1 below gives a comprehensive information on the LD_50_ values of the first ten ranked candidates.

**Table 1:** Toxicity and molecular properties of top-10 compounds.

| Compound (File Prefix) | LD<sub>50</sub> (log<sub>10</sub>(1/(mol/kg<sup>-1</sup>))) | LD<sub>50</sub> (mg/kg<sup>-1</sup>) | Molecular Weight (Da) |
| --- | --- | --- | --- |
| r1 | 2.76 | 688 | 392.05 |
| r2 | 2.12 | 1833 | 239.15 |
| r3 | 2.61 | 571 | 234.08 |
| r4 | 2.94 | 416 | 359.08 |
| r5 | 2.05 | 2371 | 266.16 |
| r6 | 3.03 | 257 | 277.13 |
| r7 | 2.41 | 915 | 235.14 |
| r8 | 2.56 | 835 | 300.15 |
| r9 | 2.68 | 560 | 265.13 |
| r10 | 2.74 | 654 | 359.18 |

Predicted LD<sub>50</sub> values (mg/kg<sup>-1</sup>) and molecular weights of selected compounds.
See Appendix C for toxicity results and comparison with GHS classification.

Chemical structure of compounds also influence drug-likeness and properties of drugs Mao et al. (2016), Jia et al. (2019). Lipinski’s rule of 5 is critical to estimate the bioavailability of a compound based on their chemical structure Benet et al. (2016), Daina et al. (2017). In figure 7, plot 1 (box plot, lipinski Violations) indicates the number of molecules with their corresponding number of violations to Lipinski’s rule. A great proportion of the molecules had zero (0/ no violations). As illustrated in plot 2 (Bioavailability Score Dist) the QED values for assessing the desirability of drugs Bickerton et al. (2012) cluster between 0.4–0.6, suggesting moderate drug-likeness, with a minority in the undesirable range (0.1–0.2).

**Figure 6:**
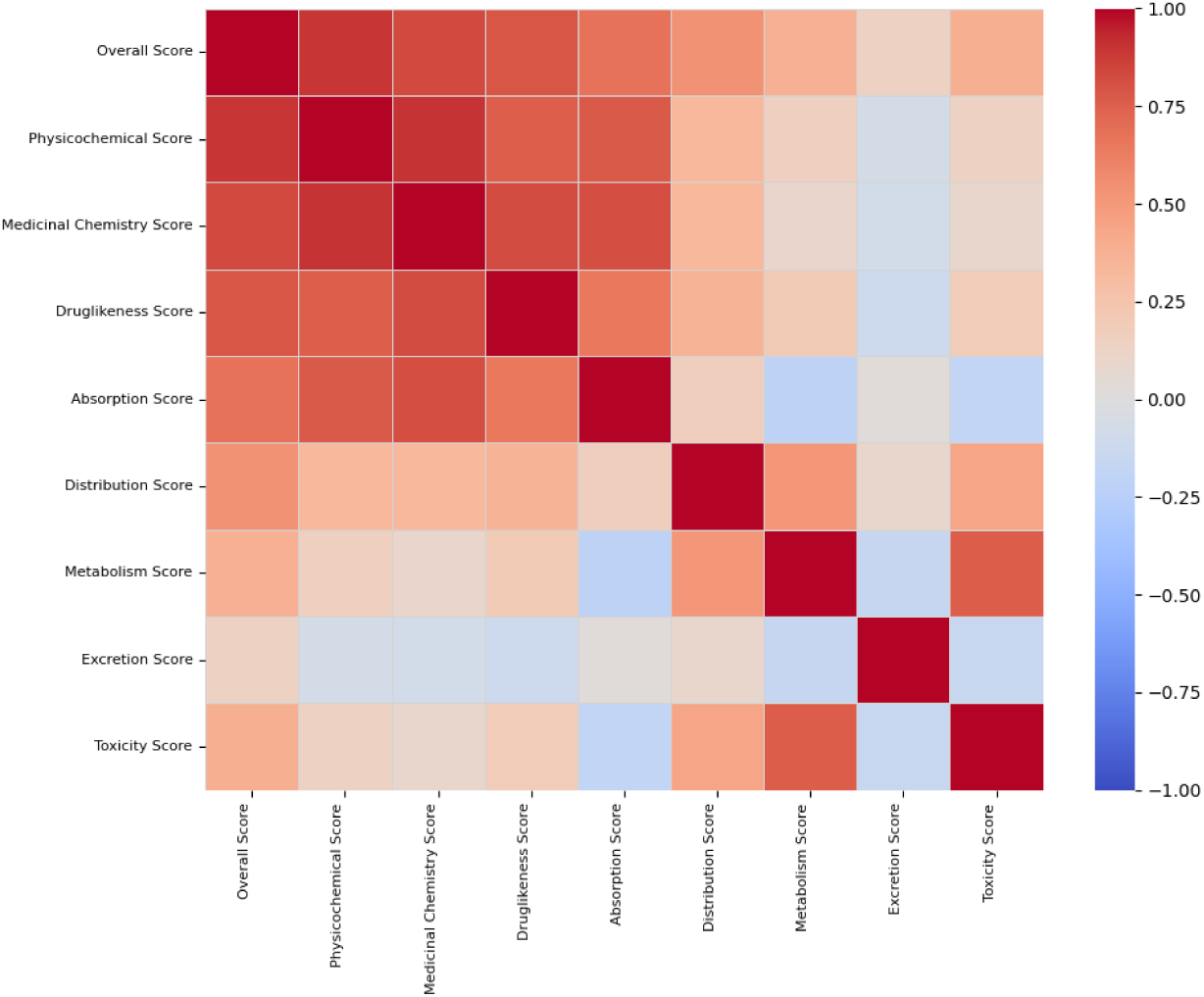
Strength of interaction between overall scores of all 8 category.

**Figure 7:**
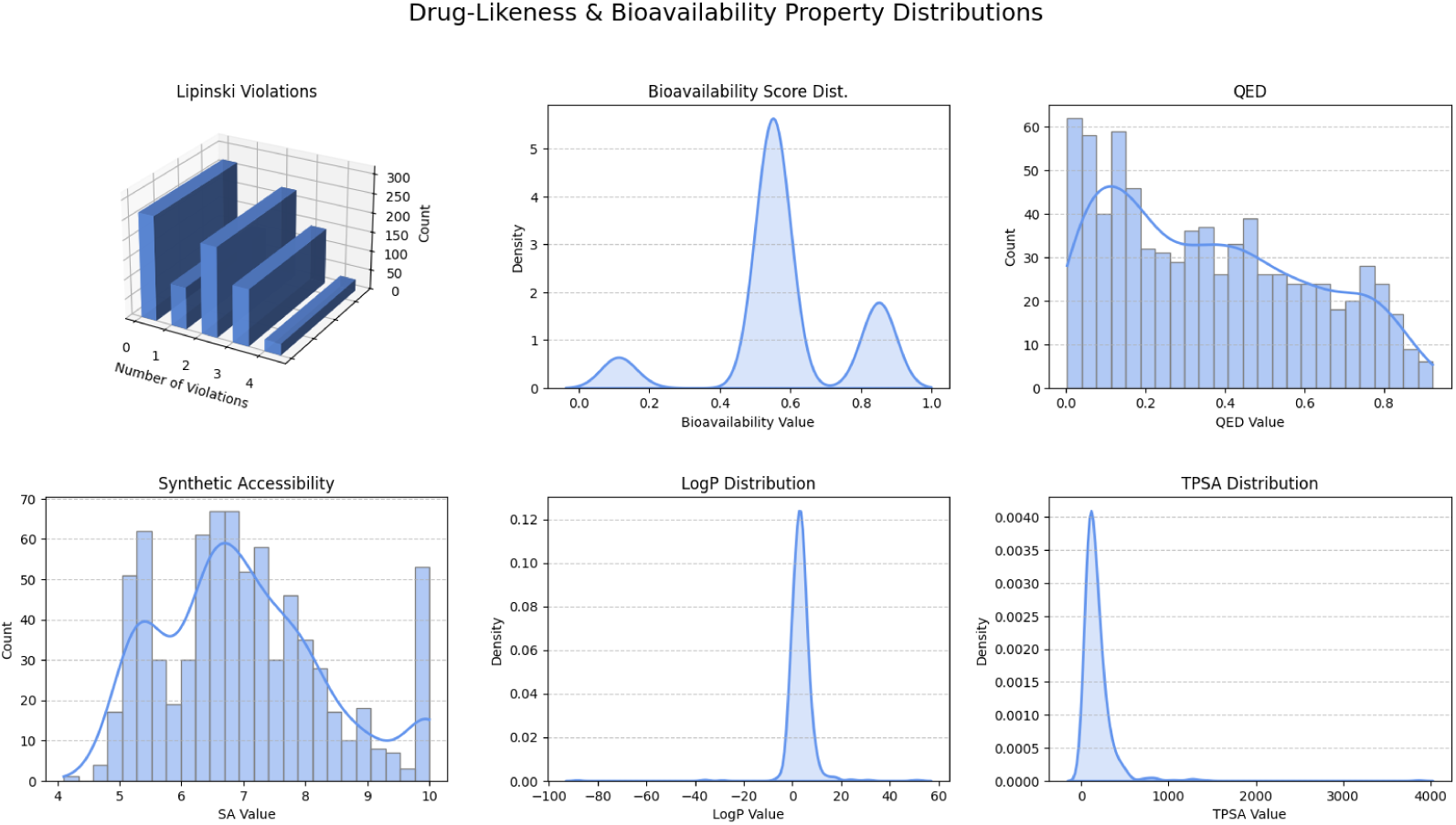
Drug-likeness and Bioavailability Property Distribution of molecules.

Subplots: **1** Lipinski violations **2** Bioavailability Property Dist 3 (Quantitative Estimate of drug-likeness) QED **4** Synthetic Accessibility **5** LogP Distribution **6** TPSA Distribution

Considering the first sixty ranked (top-60) molecules based on their overall scores, the majority fall in an acceptable range of *≥* 0.8. This threshold of *≥* 0.8 is significant because molecules achieving this score demonstrate superior ADMET properties, making them promising candidates for further experimental studies

The graph demonstrates that all 60 top-ranked molecules exhibit desirable ADMET properties and drug-likeness characteristics, with the majority achieving scores *≥* 0.8. See Appendix B for detailed ADMET predictions on top-10 molecules.

### 4.7 Antibiotics-Target Interaction

Binding Affinity prediction provides insights into the strength of interaction between molecules Kastritis and Bonvin (2013). The Binding free energy (Δ*G*) prediction using automation through Autodock VinaEberhardt et al. (2021) showed that a total of 393 molecules exhibited quantifiable bind for Fabl while 391 did for efflux pump. A total of 391 molecules exhibited binding interactions with both targets.

The computational analysis of the binding free energy (ΔG) for FabI (mean ΔG = *−*7.28 *±* 1.13 kcal/mol); and (mean ΔG = *−*7.13 *±* 0.92 kcal/mol) for AcrAB–TolC pump.

The Binding free energy (ΔG) prediction results for the top-ten performing molecules based on the ADMET ranking revealed that nine of ten of these candidates achieved moderate ΔG values against FabI, spanning –7.398kcal/mol to –5.132kcal/mol and AcrAB–TolC pump ranging from – 7.495kcal/mol and –5.505kcal/mol; this is indicative of moderate binding interactions [See Appendix D for detailed binding interaction of top-10 ranked molecules]. In contrast, for molecule rank 4 no binding free-energy estimates were predicted with both targets.

The docking poses for AnBkr1 through AnBkr10 within the FabI binding site are depicted in Figure 11.

**Figure 8:**
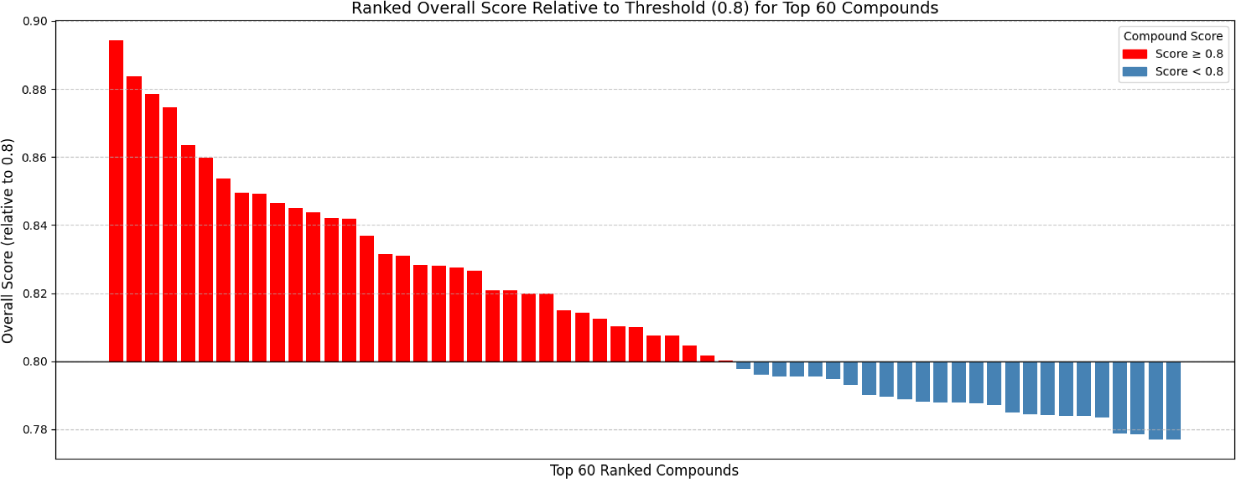
Ranked Overall Score relative to threshold (*>*= 0.8) for top 60 compounds.

**Figure 9:**
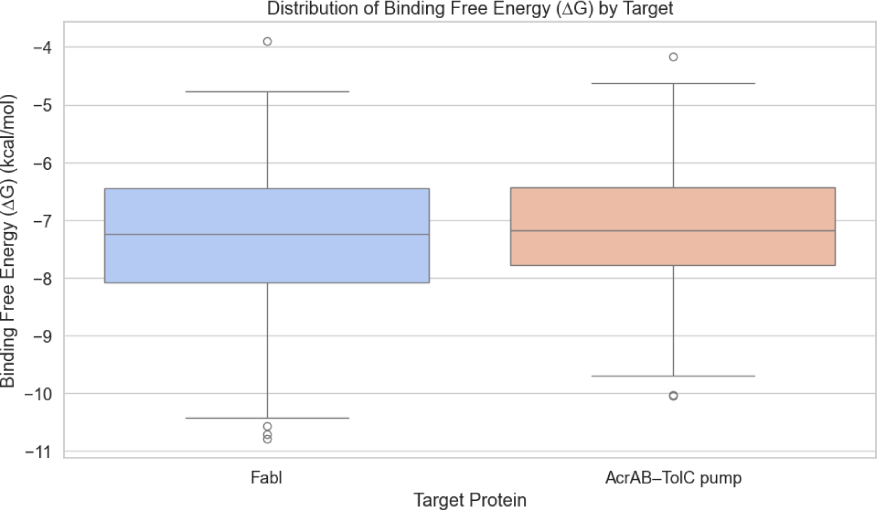
Binding free energy (ΔG) prediction.

**Figure 10:**
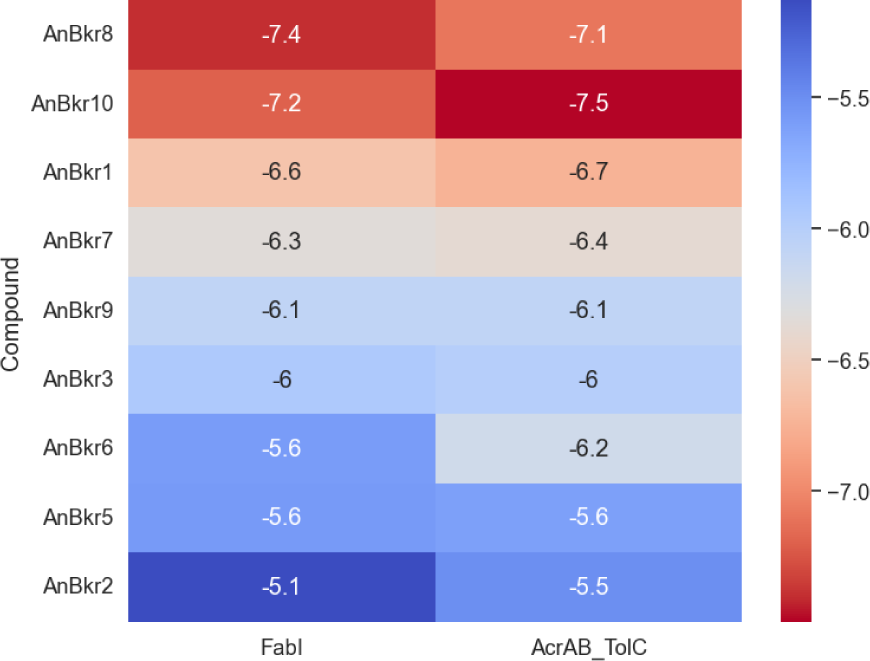
Heatmap of Binding Free Energies (kcal/mol) Across Dual Targets. Reveals Patterns of Dual Inhibition. This visualization highlights that these molecules have potential dual-target efficacy.

**Figure 11:**
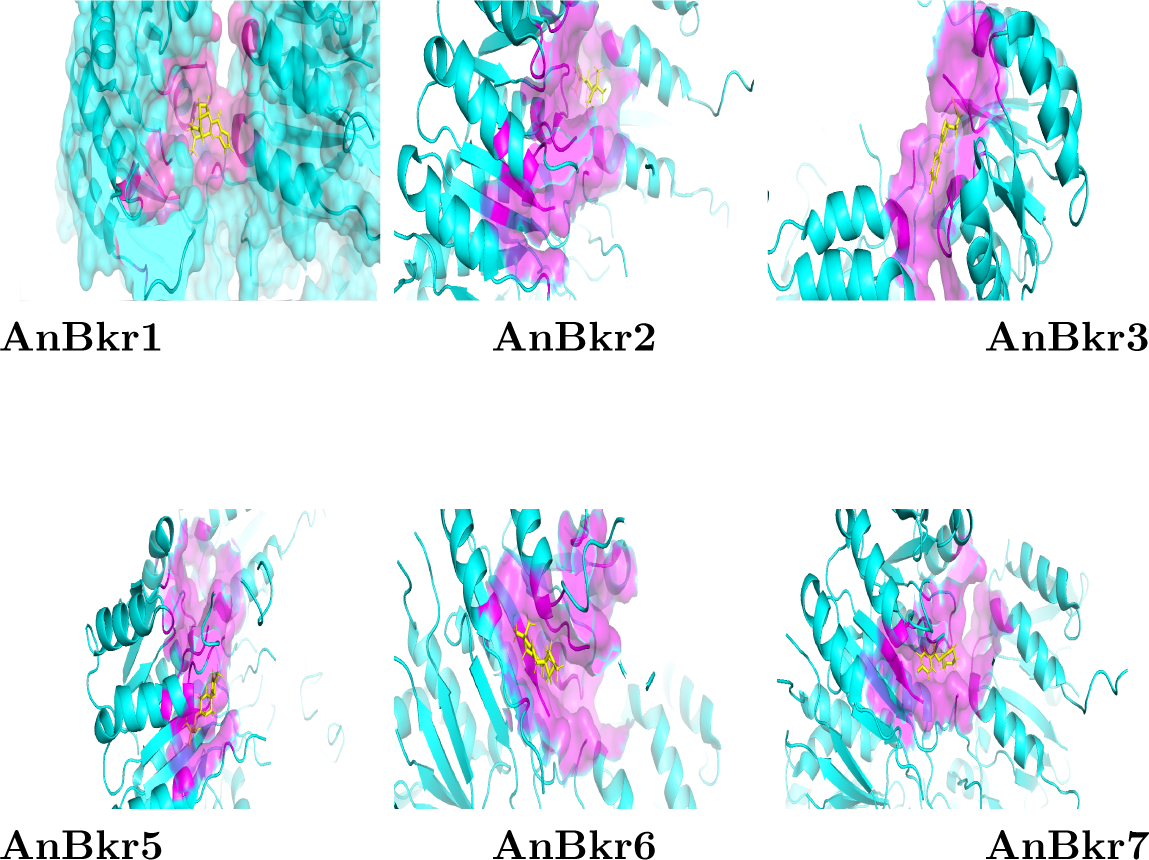
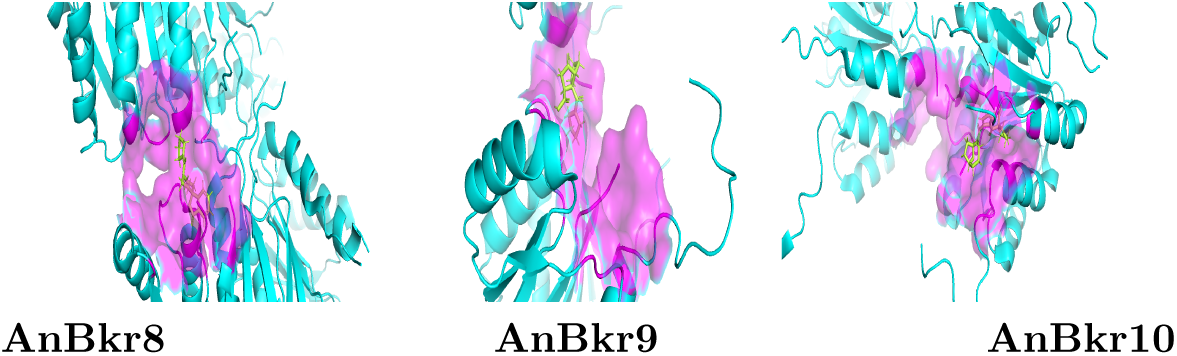
Binding interaction of top-10 molecules with the Fabl target.

Images are arranged in a 3×3 grid. The protein backbone is rendered in cyan, residues defining the binding site are highlighted in magenta, and the bound ligand is shown in limon.

More negative binding free energies (ΔG) typically indicate stronger and more favorable interactions between ligands and molecular targets, with values between –7 and –10 kcal/mol often correlating with low micromolar to nanomolar affinities. The mean binding free energy (ΔG) of the novel compounds against FabI is consistent with literature values reported for 1,3,4-thiadiazolyl thiourea derivatives, which exhibited docking scores ranging from –6.56 to –8.67 kcal/mol against FabI George et al. (2014).

Regarding the AcrAB-TolC efflux system, our molecules’ binding free energies fall within a similar energetic range to those of known inhibitors. Recent research has uncovered several pyranopyridine derivatives as potent inhibitors, exhibiting strong binding to the AcrB component of the transporter, with docking scores ranging from –10.168 to –8.427 kcal/mol Mahey et al. (2024). These findings suggest that the newly designed compounds may exhibit binding interactions comparable to known FabI and AcrB inhibitors, supporting their potential as promising candidates for further antibacterial evaluation.

## 5 Discussion

In 2021, AMR was associated with 4.71 million deaths, predominantly affecting low– and middle-income countries (LMICs) Naghavi et al. (2024). The World Health Organization has identified carbapenem-resistant Enterobacteriaceae (CRE), including K. pneumoniae, as critical AMR global health threats. The escalating prevalence of multidrug-resistant Gram-negative bacteria, particularly within the Enterobacteriaceae family, underscores the urgent need for innovative therapeutic strategies. This research proposed to use Moremi Bio Agent, an agentic LLM, to accelerate the discovery of broad-spectrum antibiotics against Enterobacteriaceae. By utilizing an agentic LLM, we aimed to rapidly design unique antibiotic molecules with dual-target capabilities, significantly reducing the time required compared to traditional methods. The integration of computational assessments for ADMET properties and binding affinities further streamlines the validation process, minimizing reliance on time-consuming procedures.

From the results shown, it indicates that our model designed molecules with promising ADMET properties. In contrast, the synthetic accessibility of molecules requires optimization; as indicated from subplot 4 (Synthetic Accessibility) there is a potential for difficulty in chemical synthesis (SA values *≥* 5) and potential high manufacturability cost for approximately 98% of generated molecules. Our results demonstrate the value of early in silico evaluations in identifying potential ADMET liabilities and prioritizing compounds for downstream experimental validation Roney and Mohd Aluwi (2024).

The results provided from this high-throughput screening of our generated molecules, have given us the ideas into which areas are necessary for optimization and molecules that can stand candidacy for further refinements and optimizations. Further research alongside refinement are required to generate promising candidates with low synthetic accessibility scores and that can interact with both targets in high affinity to further solidify our hypothesis of the dual-target mechanism of generated molecules. By harnessing this LLM technology, we aim to uncover innovative chemical scaffolds beyond the reach of traditional methods, and by pursuing a dual-target approach, we seek to overcome antibiotic resistance.

### 5.1 Limitations

All results reported here are derived solely from in silico analyses and therefore demand extensive empirical validation. To establish the true efficacy and safety profile of the generated compounds, follow-up studies must extend beyond computational predictions to include comprehensive assessment of target engagement, antimicrobial potency under physiologically relevant conditions, among others. Moreover, confirmation of any synergistic or additive effects will require functional combination assays under varying conditions to capture kinetic as well as static interactions. Finally, it should be noted that a subset of the screened molecules showed no detectable Fabl and AcrAB-TolC efflux-pump activity, underscoring the importance of further experiments to rule out false negatives and to fully characterize their resistance-modulating potential.

### 5.2 Innovation and global impact

This research addresses a critical gap in antibiotic discovery by integrating LLM approach to rapidly generate novel, dual-target antibiotic candidates against Klebsiella spp with broad-spectrum activity for Enterobacteriaceae. This innovative approach aims to overcome the limitations of traditional methods, which are often time-consuming and costly. Beyond its immediate scientific contributions, this research offers a scalable and cost-effective framework for future antibiotic discovery, ensuring a sustainable pipeline of new therapeutics to mitigate the AMR crisis. By employing advanced AI techniques, the project seeks to expedite the identification of effective antibiotics, mitigate the economic and health burdens associated with AMR and thereby contribute to global efforts in combating AMR. By focusing on AI-driven antibiotic discovery, this research not only addresses the immediate need for new antibiotics but also establishes a framework for integrating AI in drug development. This project exemplifies a pioneering effort to harness AI in addressing one of the most pressing health challenges of our time, with the potential to deliver substantial benefits on a global scale.

## 6 Conclusion

This study demonstrates the potential of AI-driven molecular design to rapidly generate and prioritize novel antibiotic candidates with promising ADMET and drug-likeness profiles. While several compounds exhibited favorable in silico properties and moderate binding to Fabl and AcrAB-TolC efflux-pump, further optimization and experimental validation are needed. The inability to identify strong binders for both targets highlight challenges in the dual-target drug design and underscores the need for improved computational methods or alternative chemotypes. Future work will focus on iterative optimization of the most promising candidates and at most possible synthesis, and in vitro/in vivo testing.

## Appendices

## A Ranking of Small Molecules

A ranking system integrated into the Moremi Bio Agent pipeline to assign ranks to molecules based on their molecular performance comprehensively. This ranking system ranks the molecules based on eight key property categories, with each category comprising multiple metrics. The table below outlines the key properties and their corresponding metrics that affect the ranking of all generated compounds.

**Table 2:**
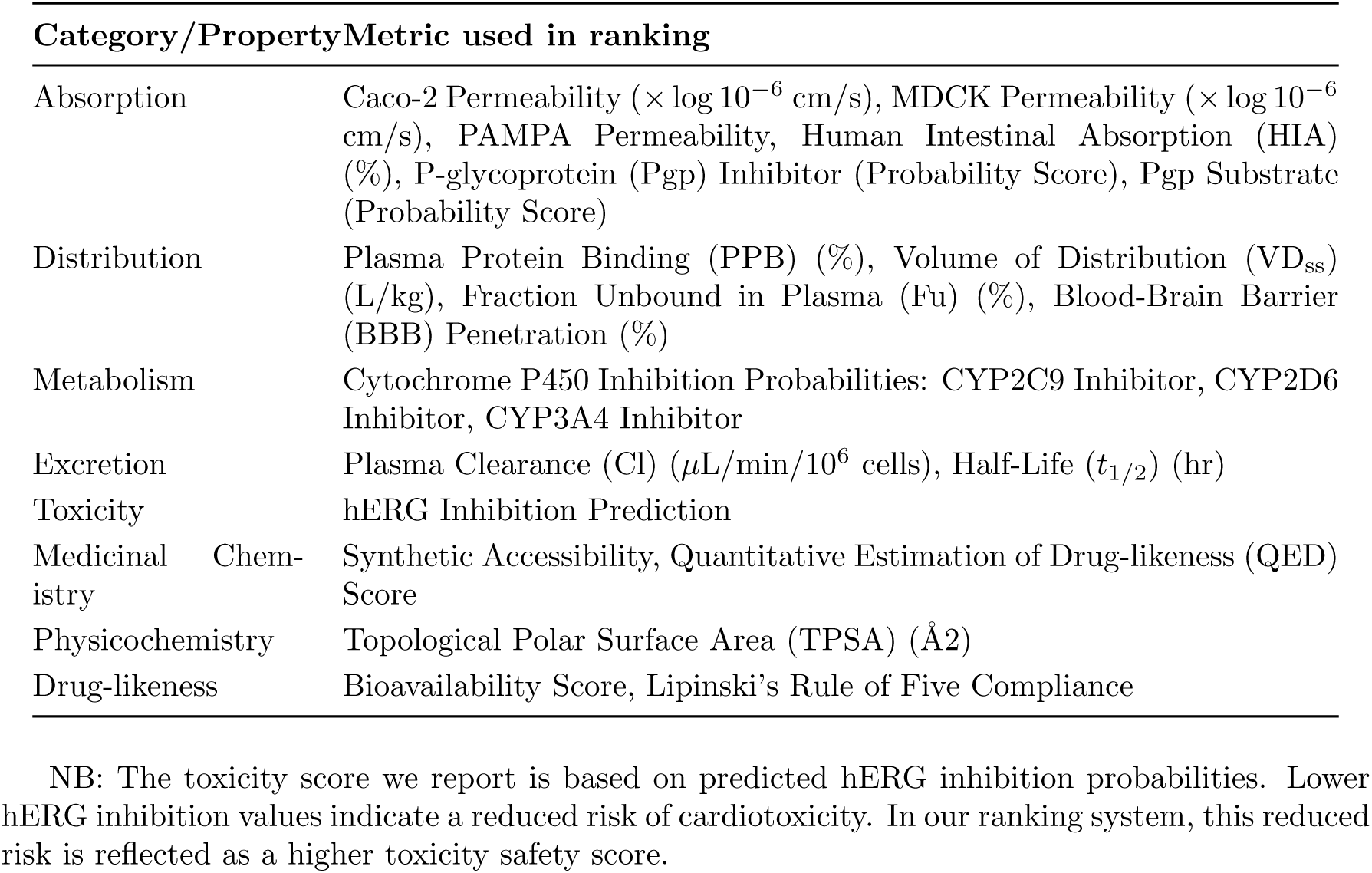
Categories and metrics used in compound ranking.

## B ADMET characteristics of top-12 molecules

The following results provide comprehensive insights into the twelve most promising candidates with broad-spectrum activity against Enterobacteriaceae based on the ranks. The visual below for top-12 promising candidates from (AnBkr1 – AnBkr12).

**Figure 12:**
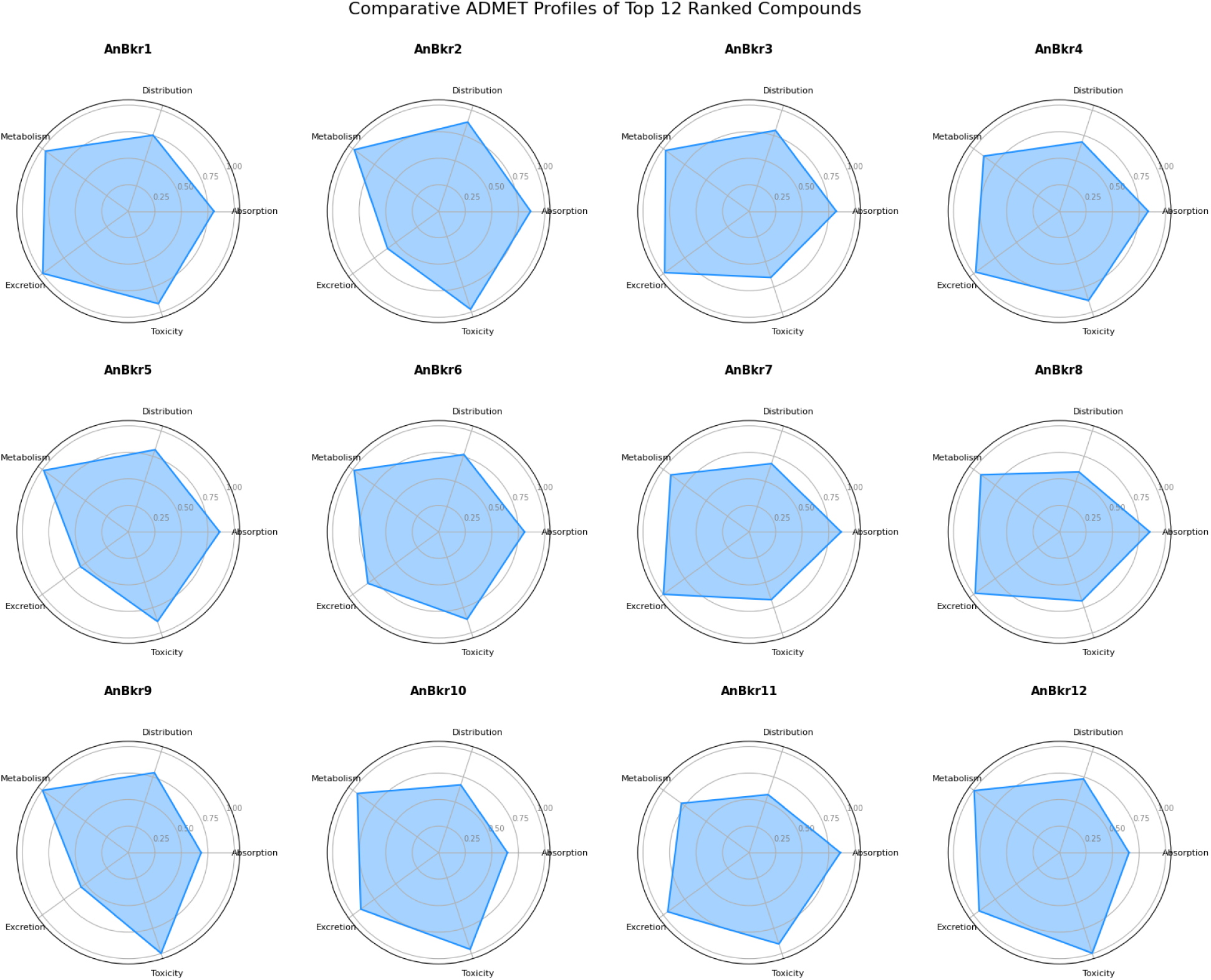
Comparative ADMET Profiles of Top-12 Ranked compounds.

## C Toxicity

LD_50_ classification based on Globally Harmonized System (GHS) for chemical classification for top-10 ranked molecules.^2,3^

**Table 3:** Distribution of molecules across toxicity categories based on LD_50_ values.

| Compound (r) | LD<sub>50</sub> (mg/kg) | Toxicity Category |
| --- | --- | --- |
| AnBkr1 | 688 | Category 4 |
| AnBkr2 | 1833 | Category 4 |
| AnBkr3 | 571 | Category 4 |
| AnBkr4 | 416 | Category 4 |
| AnBkr5 | 2371 | Category 5 |
| AnBkr6 | 257 | Category 3 |
| AnBkr7 | 915 | Category 4 |
| AnBkr8 | 835 | Category 4 |
| AnBkr9 | 560 | Category 4 |
| AnBkr10 | 654 | Category 4 |

Distribution of Molecules Across Toxicity Categories Based on LD<sub>50</sub> (mg/kg) Values

Table 4 summarizes the distribution of predicted LD_50_ values for the generated antibiotic candidates. A majority of the compounds (51.16%) fall under Category 4 (Harmful), indicating moderate toxicity levels. Notably, 23.77% of the molecules are classified as Category 3 (Toxic), suggesting a need for structural optimization in this subset. Conversely, 13.05% of the candidates were found to be practically non-toxic, with LD_50_ values exceeding 5000 mg/kg, while 10.85% exhibit low toxicity (Category 5). A small fraction (1.16%) demonstrated high toxicity (Category 2), representing high-priority targets for exclusion or redesign. These findings provide an initial safety profile landscape for the antibiotic library and guide further selection for in vivo validation.

**Table 4:**
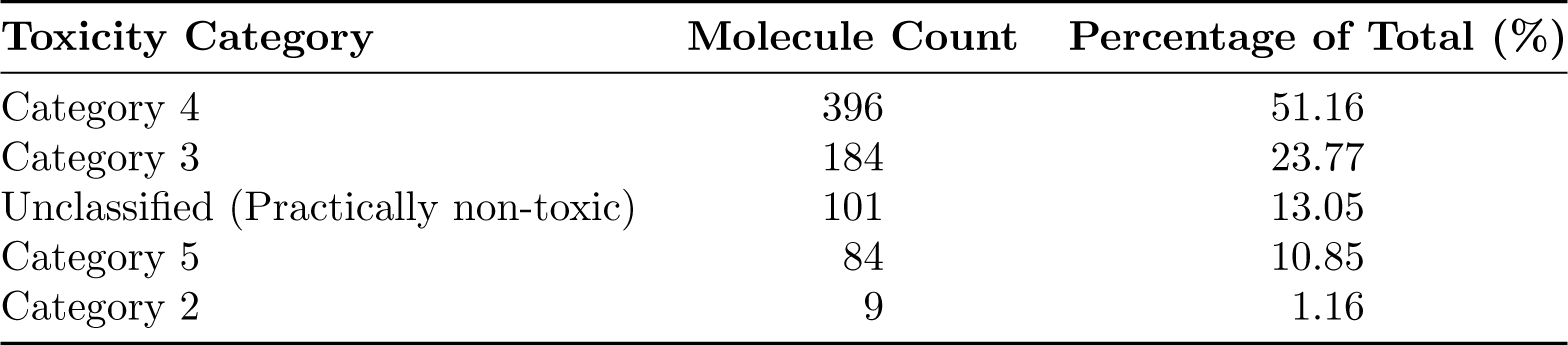
Distribution of molecules across toxicity categories.

## D Binding Free Energy and Potential Dual-target Inhibition of Top-10 Ranked Molecules

The figures below present the comparative binding interactions of top-10 candidate compounds across dual targets. Bar chart showing the binding free energies (kcal/mol) of computationally generated antibiotic candidates against two bacterial targets. Each compound is evaluated across both targets to assess dual-inhibition potential.

**Figure 13:**
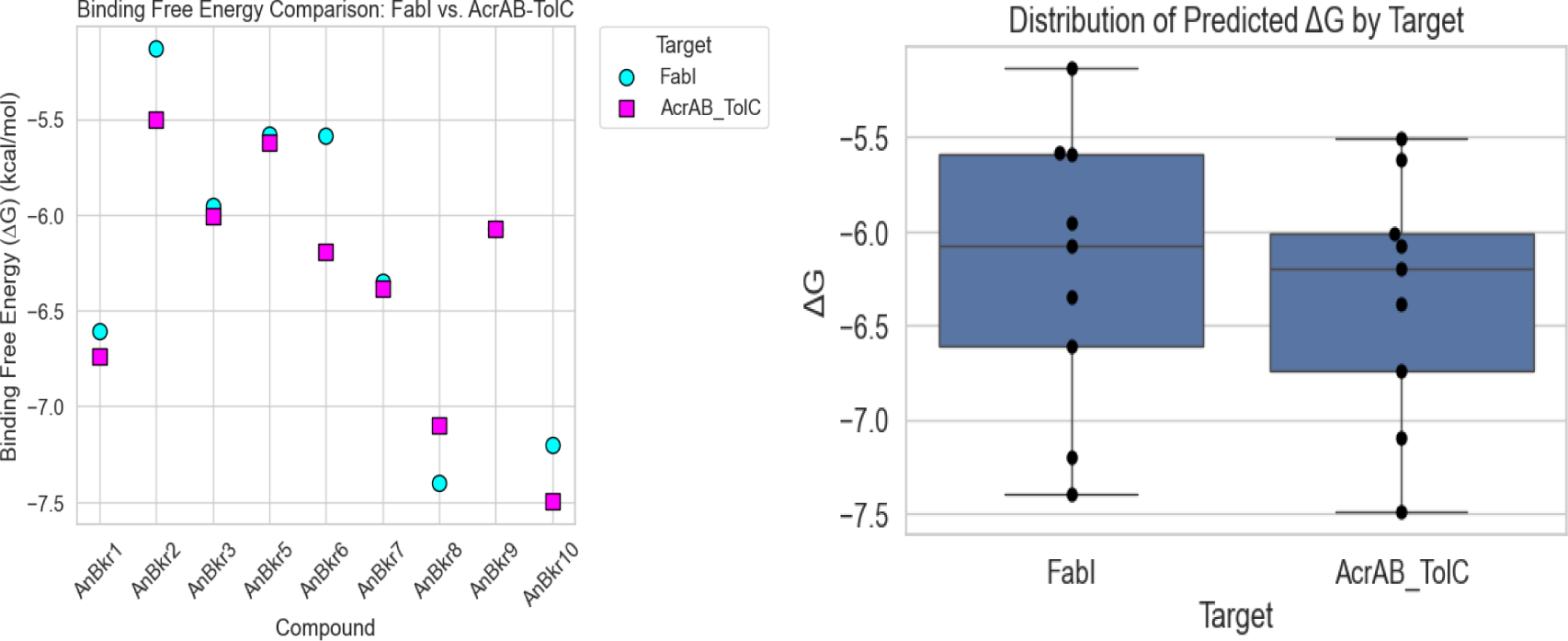
Distribution comparison across ADMET properties.

The binding free energies (kcal/mol) of the top 10 candidate compounds on dual targets.

**Table 5:** Binding free energies ΔG of top-10 ranked molecules.

| Molecule Rank | Binding Affinities (kcal/mol) |  |
| --- | --- | --- |
|  | FabI | AcrAB-TolC efflux-pump |
| AnBkr1 | -6.61 | -6.74 |
| AnBkr2 | -5.132 | -5.505 |
| AnBkr3 | -5.952 | -6.008 |
| AnBkr5 | -5.582 | -5.621 |
| AnBkr6 | -5.587 | -6.194 |
| AnBkr7 | -6.348 | -6.386 |
| AnBkr8 | -7.398 | -7.098 |
| AnBkr9 | -6.075 | -6.074 |
| AnBkr10 | -7.199 | -7.495 |

## Footnotes

1 https://gcgh.grandchallenges.org/challenge/innovations-gram-negative-antibiotic-discovery

2 https://www.ncbi.nlm.nih.gov/books/NBK253967/

3 https://webapps.ilo.org/static/english/protection/safework/ghs/ghsfinal/ghsc05.pdf?

